# Unique mutational changes in SARS-CoV-2 genome: A case study for the largest state of India

**DOI:** 10.1101/2020.08.24.265827

**Authors:** Priti Prasad, Shantanu Prakash, Kishan Sahu, Babita Singh, Suruchi Shukla, Hricha Mishra, Danish Nasar Khan, Om Prakash, MLB Bhatt, SK Barik, Mehar H. Asif, Samir V. Sawant, Amita Jain, Sumit Kr. Bag

## Abstract

COVID-19 is a global pandemic causing more than 8 million deaths till mid-August, 2020. In India, more than 3 million confirmed cases have been reported although with relatively low death rate of 1.8%. In this study, we sequenced 47 genomes of SARS-CoV-2 from the patients of 13 districts of Uttar Pradesh (UP), the largest state of India using third-generation sequencing technique. The phylogenetic clustering revealed that no UP sample was aligned with the previously defined USA clade, where the mortality was high. We identified 56 distinct SNP variations in the genomes of UP resulting in a unique mutation rate of 1.19% per sequence, which is greater than the value 0.88% obtained for the rest of India. The relatively less death rate in UP indicates that the mutation in the virus is deleterious. Further investigation is required with larger sample size to determine the degree of virulence vis-a-vis SNP variation.

## 1. Introduction

COVID-19, a global pandemic [1] has been the focus of many researchers worldwide to identify the potential region(s) of its pathogenicity since its emergence in December 2019 [2]. The causative agent, SARS-CoV-2 belongs to the Coronaviridae family and comprises of single-stranded positive-sense RNA with a genome size of approximately 30 kb [3]. SARS-CoV-2 genome consists of 29 open reading frames (ORFs), among which four are structural proteins (envelope protein, membrane protein, nucleocapsid protein, and spike protein), some accessory proteins translated by ORF3a, ORF6, ORF7a, ORF7b, ORF8, and ORF10, and 16 nonstructural proteins (nsps) [4]. During the spread of COVID-19, the virus evolved continuously resulting in variation in death rates in different countries [5]. As of 20^th^ August 2020, approximately 23 million confirmed cases were reported worldwide with ∼800 thousand registered deaths. At present, India ranks 3^rd^ in the total number of COVID-19 cases. However, fatality is 1.8 %, which is considered as one of the lowest death rates in the world [6] [7]. Many reports correlate the fatality to the mutational changes in the virus genome in different geographic regions [8][9,10][11]. D614G substitution at the spike protein has been related to the cause of high mortality in humans [12]. The authors of a study on the western Indian populations have correlated the mutational changes to specific age groups and modes of infection [13]. To assess mutation rates and understand the evolution of the SARS-CoV-2 genome in different continents, many genomes were sequenced by employing different techniques and submitted to GISAID [14]. More than 84 thousand full-length SARS-CoV-2 genome sequences including >2400 viral genomes from India were available on the GISAID platform as on 12^th^ August, 2020. Using this data, the Council of Scientific Industrial Research-Institute of Genomics and Integrative Biology (CSIR-IGIB) recently delineated an India-specific clade [15].

Currently, out of 29 states and 7 Union Territories of India, UP ranks 5^th^ in confirmed number of cases of COVID-19. As on 12^th^ August, 2020, only 11 sequences were available on the GISAID database for UP that represented only a small population in four districts. Considering the extremely low number of genomes in the database despite high number of infected persons in the state, it is required to sequence many more genomes of the virus from UP. It is also essential to understand the mutation pattern(s) causing variation in genome, and to compare that with the national/global mutation variations so that the changes that affect the properties of virus e.g. by making some lineages more or less virulent or transmissible can be established [16]. Therefore, we report 47 genome sequences of SARS-CoV-2 virus representing 13 districts of Uttar Pradesh (UP) using third-generation sequencing technique (Oxford Nanopore PromethION) in this study. The phylogenetic clustering was undertaken to classify UP genomes into different clades. We investigated the unique mutational features and rates during the transmission of the virus. We also correlated the death rates in different states including UP with their synonymous to nonsynonymous unique mutation ratio.

## 2. Methods

### 2.1 RNA Preparation and Sequencing of the viral genome

Total viral nucleic acid was extracted from the clinical sample of SARS-CoV-2 using PureLink DNA/RNA mini kit (Invitrogen, USA) as per manufacturer’s instructions as well as the guidelines of Indian Council of Medical Research (ICMR) [17]. The Human RnaseP gene was tested in all samples to check the quality of samples and to validate the process of nucleic acid extraction. Real-time RT-PCR assays were performed using the Superscript III One-Step Real-Time PCR kit (ThermoFisher Scientific, USA) as per the protocol[18]. All samples were tested by two stage protocol consisting of screening and confirmation. E-gene and RnaseP were used for screening while RdRp and ORF1b were used as confirmatory assay for SARS-CoV-2. All the samples positive for E-gene, RdRp, and ORF1ab with Ct ≤ 25 were further taken for whole-genome sequencing.

Virus genomes were generated by using ARTIC COVID-19 multiplex PCR primers followed by nanopore sequencing on two ONT PromethION flow cell[19]. For PromethION sequencing, libraries were prepared using the ligation sequencing kit (SQK-LSK109) and Native barcoding (EXP-NBD 104 and EXP-NBD 114) of 24 samples in a single flow cell. The selected samples were converted to the first-strand cDNA using reverse transcriptase as suggested in Nanopore protocol “PCR tiling of COVID-19 virus” (Version: PTC_9096_v109_revD_06Feb2020). The multiplex PCR was performed with two pooled primer mixture as recommended and the cDNA was amplified. After 35 rounds of amplification, the PCR products were collected, purified with AMPure XP beads (Beckman coulter), and quantified using Qubit4 Fluorometer (Invitrogen). The double-stranded cDNA was further subjected to end-repair, A-tailing, and Native barcode ligation individually. The barcoded samples were pooled together and ligated with sequencing adapters followed by loading of 50ng of pooled barcoded material and sequencing on two PromethION flow cells having 24 samples each.

### 2.2 Genome Assembly

Quality passed nanopore reads were subjected to remove the chimeric reads between the 300 to 500 base pair length according to the amplicon size. Using Artic network pipeline [20], we individually assembled the high coverage (∼9000X) filtered long reads of each sample. During the assembly “--normalize” option was used to normalize the data up to 200 bp coverage. This normalization is useful for the minimization of the computational requirements of our high coverage data. The SARS-CoV-2 reference genome (MN908947.3) [21] was used for generating the consensus sequence from the assembled reads. Low complexity or primer binding regions were masked during the consensus sequence generation and variant calling, to avoid the ambiguity sequences.

### 2.3 Data Preparation

For comparative and explanatory analyses of our assembled genome from UP, we downloaded the whole genome data of all Indian sequences from the GISAID database that was available until the 12^th^ of August, 2020. We randomly selected 100 sequences each from Asia, Europe, South America, North America, Central America and Eurasia as an outlier for different analyses of Indian sequences. A total of 2923 processed sequences (2323 from India and 600 from different continents) were further screened based on unrecognized nucleotides (Ns) with the full length of genome. We kept only those sequences whose unrecognized base pair is less than 1% of the total length followed by the masking of the low complexity region. Along with it, we also considered the 47 assembled genome sequences of Uttar Pradesh of India for further analysis.

### 2.4 Phylodynamics clustering of Indian sample

The phylodynamic clustering of 47 genome sequences of UP was conducted, on the basis of nextstrain defined clade [22]. For a comprehensive study of the clade, we took 856 samples of the SARS-CoV-2 genome of India that was available till 10^th^ June. Whole-genome alignment has been carried out using mafft aligner [23] by taking the Wuhan Hu-1(MN908947.3) sample as a reference genome. IQTREE software [24] was used for the phylogenetic tree with the GTR substitution model. The refinement of the tree was done by the “--timetree” parameters to adjust the branch lengths and filter out the interquartile ranges from regression. The refined time-resolved tree was further screened to derive the ancestral traits, infer nucleotide mutation as well as amino acid variation from the root. The resulting phylodynamics tree was then used to define the clade based on a new classification that signifies the geographic spreading of SARS-CoV-2 in India. The final tree was further visualized through the nextstrain shell by using the auspice interface [25].

### 2.5 Divergence estimates of the newly assembled genome

Divergence estimation of the newly assembled genome and the most recent common ancestor (tMRCA) were examined using the BEAST v1.10.4 program [26]. Through the MRCA analysis, the origin of the ancestral virus was identified. With the strict molecular clock and the exponential growth model, we processed the Markov chain Monte Carlo algorithm on the masked alignment file for 100 million steps, where the first 30 million steps were used for burn-in to get the effective sample size. Bayesian coalescent analysis of the UP samples was conducted using the GTR substitution model with the kappa scale of 1 to 1.25. Molecular clock and root divergence time were reconfirmed with the treetime [27] using the same substitution model.

### 2.6 Variant identification and functional evaluation

We scrutinized the unique mutations in our assembled genome of UP state that was called during the assembly. To identify the SNP variants among all the processed (downloaded and assembled) sequences and to ensure the uniqueness of variants in our assembled data, we aligned each downloaded sequences separately to the reference genome (MN908947.3) by use of mafft aligner [23]. The variation was detected with the snp-sites program and [28] nextstrain. The resulting Variant Call Format (VCF) file was annotated further to find out the position of the mutation that affects the SARS-CoV-2 virus genome though snpEff program [29]. Finally, manual screening of mutation was carried out to define the unique mutation in our processed sequences along with the other continents of the world. Functional validation of the emerging new mutations in UP sequences was undertaken through the Sorting Intolerant from Tolerant (SIFT) [30]. SIFT evaluated the functional consequences of the variants, where the SIFT score of 0.0 to 0.5 indicates deleterious effect and >0.05 was interpreted as a toleratable mutation. The Identified variants of UP sequences were visualized through the R package [31].

## 3. Results and Discussion

### 3.1 Demographic Representation

In this study, a total of 47 genomes of SARS-CoV-2 were sequenced using the long reads PromethION sequencer. All the samples were from UP state of India representing 13 different districts (**Fig 1a**). The samples were collected from symptomatic as well as asymptomatic patients in which the male-female ratio of patients was 3:1. The age of the patients varied between 2 and 86 years with a mean age of 35 years. The maximum number of samples were collected in May, 2020. Only the sample numbers 2 and 13 were collected in the months of March and April, 2020, respectively (**Supplementary File 1**). The mean coverage of all the samples was 9000X which was related to the ct value of the samples (**Fig. 1b**).

**Fig. 1.**
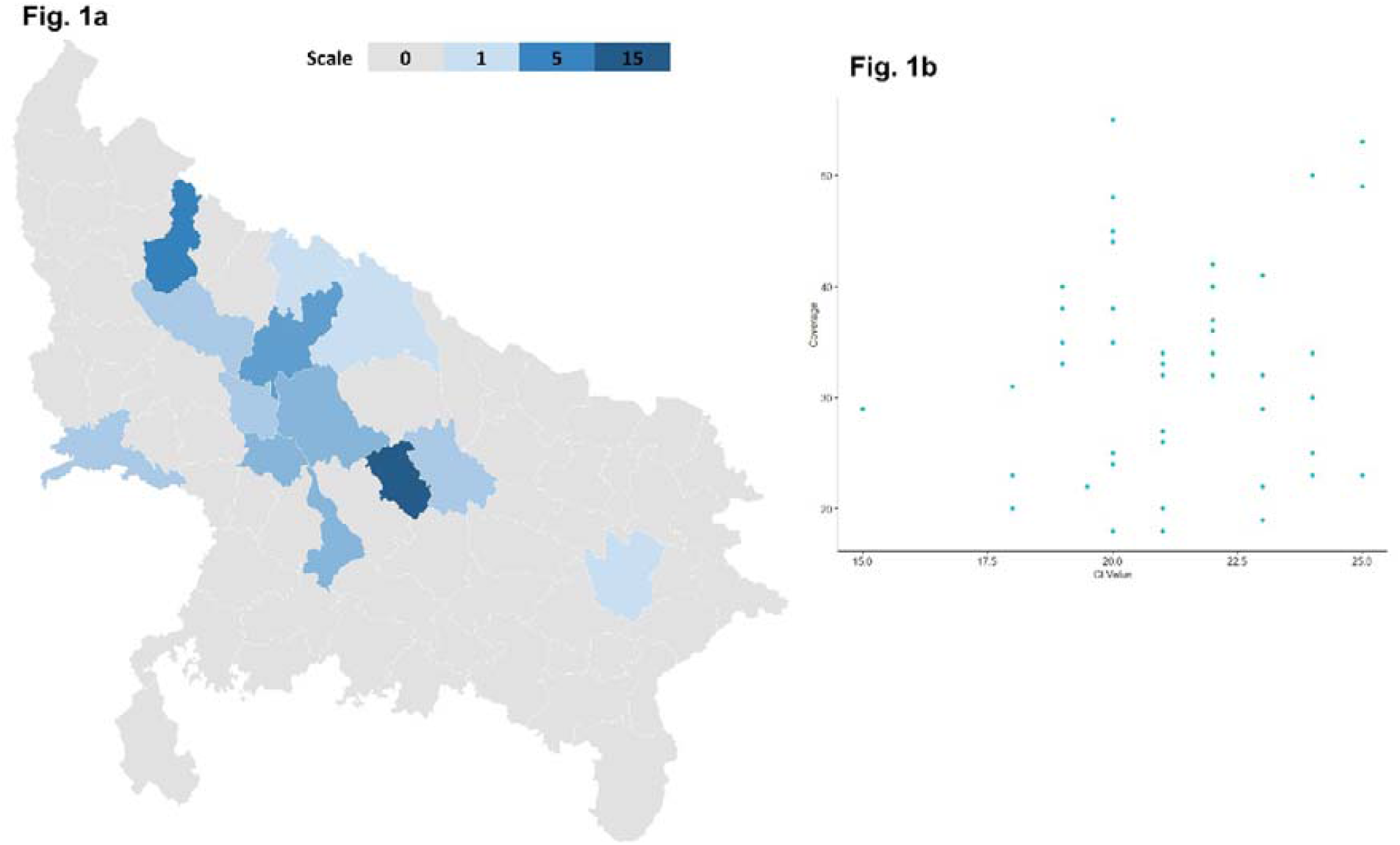
a) The detailed map of the SARS-CoV-2 genome sequences of different districts of the Uttar Pradesh state of India. The scale denoted the number of sequenced genome. Grey color means no genome was sequenced while the darker blue color represents the highest number of genome sequencing. b) The Co-linear plot of the Ct value of 47 positive COVID-19 patients to the coverage of the whole genome sequence long-read data. Each sample was represented through the blue dots where higher Ct value shows high coverage data.

### 3.2 Assembly of the whole genome

The resulting fastq and fast5 files of the nanopore PromethION sequencer were quality checked by the artic network pipeline [20]. The amplicon reads of 300 to 700 base pair length were removed from the raw fastq file to avoid the generation of chimeric reads. Filtered fastq file and raw fast5 format files of each sample were taken as input for whole-genome assembly. The MN908947.3 genome was used as a reference to generate the consensus sequence for each sample individually. Additionally, medaka parameters were also used to identify the variants or SNPs based on assembled reads during the generation of the consensus sequence. To avoid the ambiguity at the time of SNP calling, the primer binding region in the genome was masked. The resulting Single-nucleotide Polymorphism (SNPs) were considered as true SNPs in our sequenced samples data because they had high-quality read depth (∼300) at each base pair. These High-quality SNP reads were used to define the unique mutation features in the SARS-CoV-2 genome sequences of UP in comparison to the sequences from other states of India as well as from different continents. The assembled whole-genome sequences were submitted to the GISAID with accession ID (EPI_ISL_516940-EPI_ISL_516986). The metadata information is available in **Supplementary File 1**.

### 3.3 Phylodynamic analyses of the assembled genome

Phylodynamic clustering was used for clade assignment in our assembled genome based on nextstrain where we could trace the origin of the clade concerning the specific mutation on the genomic region. A total of 47 assembled genomes of SARS-CoV-2 from 13 different districts of UP in India, and 856 genomes from 21 different states of India as available on 10^th^ June were used to perform the phylodynamic clustering according to the new clade classification by taking the Wuhan SARS-CoV-2 reference genome (EPI_ISL_406798) as an outlier. The information on the sequences including their accession number, sample name, collection date is mentioned in **Supplementary File 2**.

The augur pipeline [32] was used for the alignment of all the sequences, their phylogenetic clustering, defining the interquartile range and deciphering the time resolving tree, using SARS-CoV-2 (MN908947.3) genome as the root. The resulting filtered tree was subjected to define the potential emerging clades based on nucleotide mutation(s). Indian samples were clustered into earlier designated five clusters delineated by Nextstrain viz., 19A, 19B, 20A, 20 B, 20C (**Fig. 2a**) [33]. According to these authors, two clades viz., 19A and 19B were primitive and dominated by Asian countries, and were delineated by C to T mutation at 8782 position in ORF1ab region, and T to C mutation at 28144 position, respectively. A total of 43% of processed sequences of India (380 out of the 880) were aligned with these two clades.

**Fig. 2.**
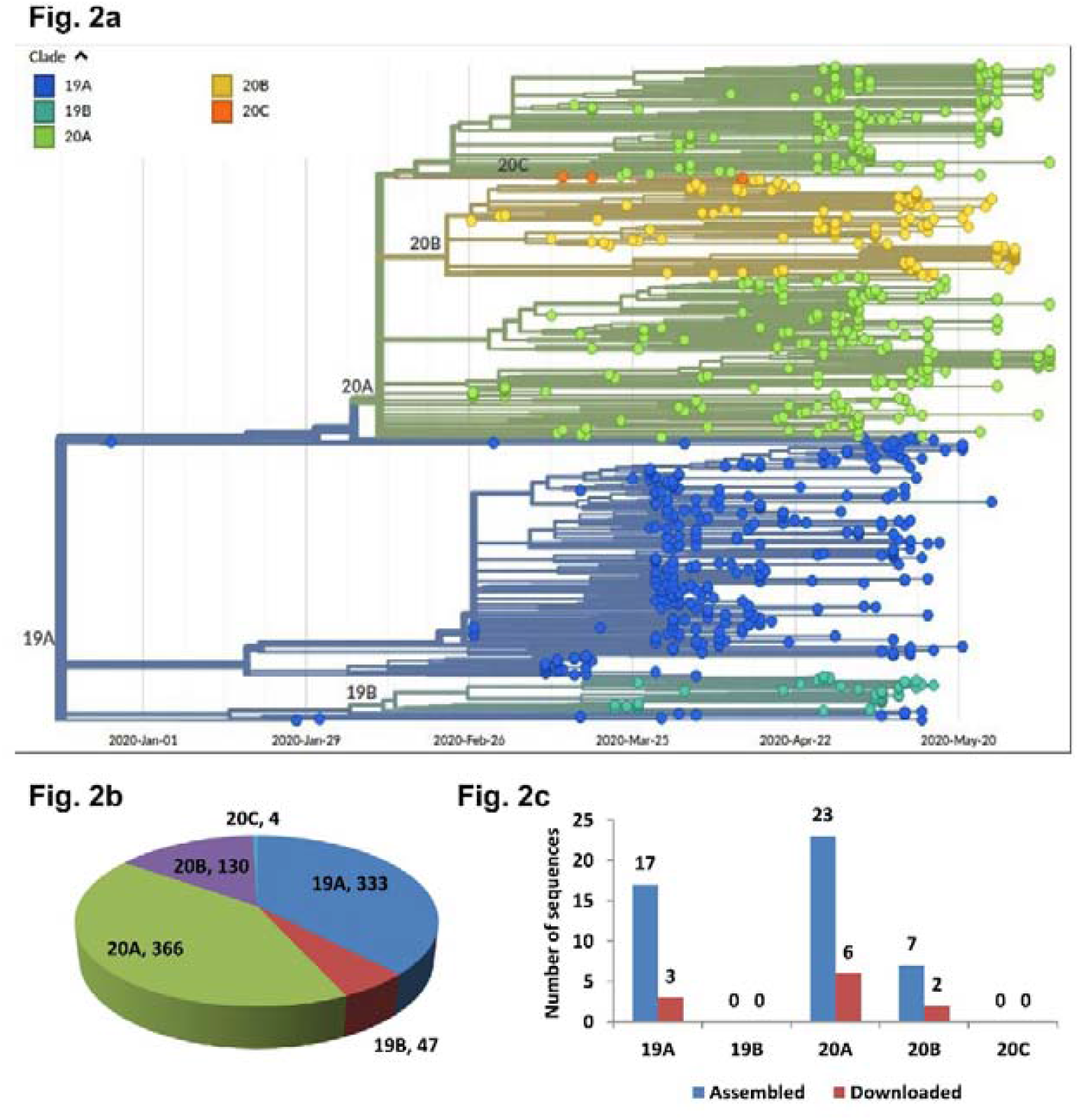
a) The phylogenetic clustering of 880 samples of India according to the nextstrain defined clade. The 19A and 19B clades are the primitive clades that were sequenced in early 2020; whereas 20A, 20B and 20C clades are designated as emerging clades. The reference genome of the Wuhan sample was implemented in the clustering trees which aligned in 19A clade before the starting of the Jan-2020. b) Distribution of sequenced virus genome of India on different clades that were visualized through the piechart. C) Classification of Uttar Pradesh SARS-CoV-2 genome on different clades. The blue color bar shows the 47 assembled sequences from this study and red color bar represents 11 downloaded sequences from GISAID for UP.

The clades 20A and 20B were considered as European clade by these authors based on the dominance of European sequences in these two clades. The clade 20C is well known as North American cluster [33]. The remaining Indian sequences (out of 880 total) were aligned with these three clades, which deviated from the root (19A) in early 2020. Most of the Indian sequences (366) were clustered into 20A clade, followed by 19A clade (333) (**Fig. 2b**). The least number of Indian sequences belongs to 20C clade. The United State of America (USA) is the primary source for 20C clade and only 4 Indian sequences (two each from Delhi and Gujarat) were aligned to it. The sequences from Delhi (EPI_ISL_435063, EPI_ISL_435064) did not show any nucleotide or amino-acid mutation, while sequences from Gujarat (EPI_ISL_435051, EPI_ISL_435052) had two missense variations (A55C, C241T) in ORF1ab region.

Out of 47 SARS-CoV-2 genome sequences obtained in the current study for UP, 17 sequences belong to 19A clade, 23 to 20A and 7 to 20C clade. None of the sequences were aligned with 19B and 20C clades **(Fig. 2c)**. Eleven sequences for UP, submitted by earlier workers to the Nextstrain also belong only to 19A, 20A and 20C clades. Although originally nextstrain global page classified these two clades (19B and 20C) as Asian and North American based on the mutation pattern and the respective continental dominance of sequences, the increasing number of sequence deposition to the database is now somewhat exhibiting a mixed geographical distribution.

### 3.4 Estimation of the date of origin

The mutation rate of the 47 sequenced genomes was estimated through BEAST software using the coalescent model as the prior of the trees [26]. The estimated rate of mutation was predicted as 3.279e-04 substitutions per site per year for the given SARS-CoV-2 genome sequences. Highest Posterior Density (95 %) intervals are in the range of 2.24e-04 to 4.459e-04. (**Supplementary File 3**). The substitution rate was further confirmed through the treetime software [27] where root to tip regression rate was calculated as 3.02e-04 with the 0.05 correlation value (**Fig. 3**). This correlation value indicates the informative behavior of inputs for temporal information to rationalize the molecular clock approach. The root date of the SARS-CoV-2 genome from UP state was estimated as 29 December, 2019 which substantiated the reported timing of this virus in Wuhan city of China [34].

**Fig. 3.**
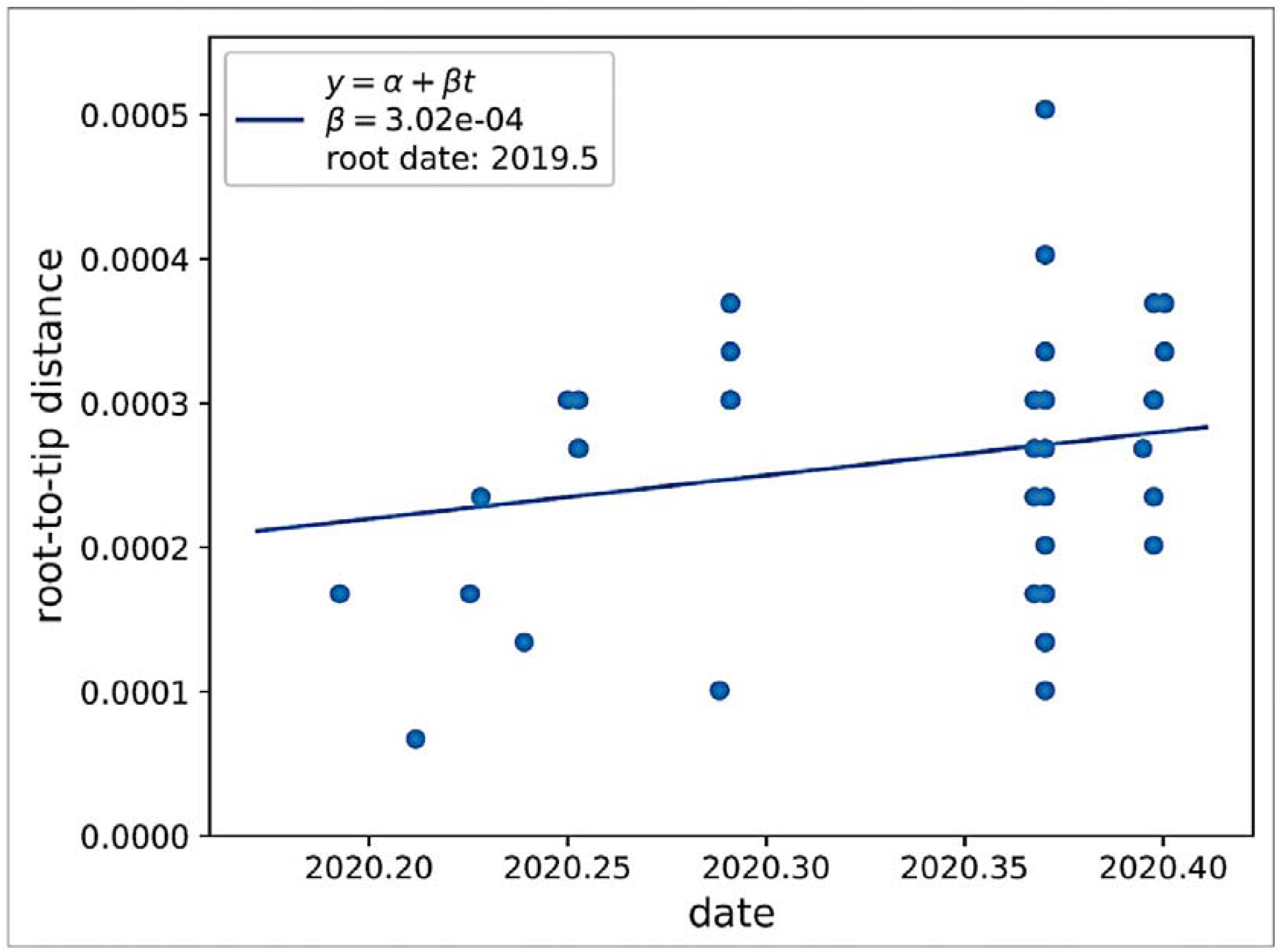
The root-to-tip regression rate of sequenced SARS-CoV-2 genome of Uttar Pradesh state of India to investigate the origin of infection. X-axis describes the timing of the sample collection of COVID-19 infected persons. Y-axis describes the regression rate in comparison to the sample collection date. The origin of infection in Uttar Pradesh samples is estimated by 2019.5.

### 3.5 Assessment of unique Mutational features in different states of India

We identified the unique mutational features in the SARS-CoV-2 genomes of other states of India. The unique mutation rate is ∼0.88% of the total sequenced genomes with 0.42 synonymous to nonsynonymous mutation ratio. For the isolation of unique SNP features, we selected only those states for which at least 10 sequences of SARS-CoV-2 genome were available as on 10^th^ of August, 2020. Thus, 12 states viz., Gujarat, Maharashtra, Telangana, Delhi, Odisha, Karnataka, West Bengal, Uttarakhand, Madhya Pradesh, Tamil Nadu, Haryana and Uttar Pradesh were selected in order of descending sequenced genome numbers **(Supplementary File 4)**. We strike off the common mutation positions from different continents (Asia, Africa, Eurasia, North America, Oceania, and South America) to check the mutated version of the SARS-CoV-2 genome in the Indian strain. The highest number of unique mutation was annotated in Maharashtra followed by Gujarat, Odisha, Telangana, West Bengal and Karnataka **(Table 1)**. The unique SNP information was annotated on the SARS-CoV-2 reference genome (MN908947.3) and classified it into synonymous, missense, stop gain, stop loss, upstream gene variant and downstream gene variant categories **(Fig. 4)**.

**Table 1.**
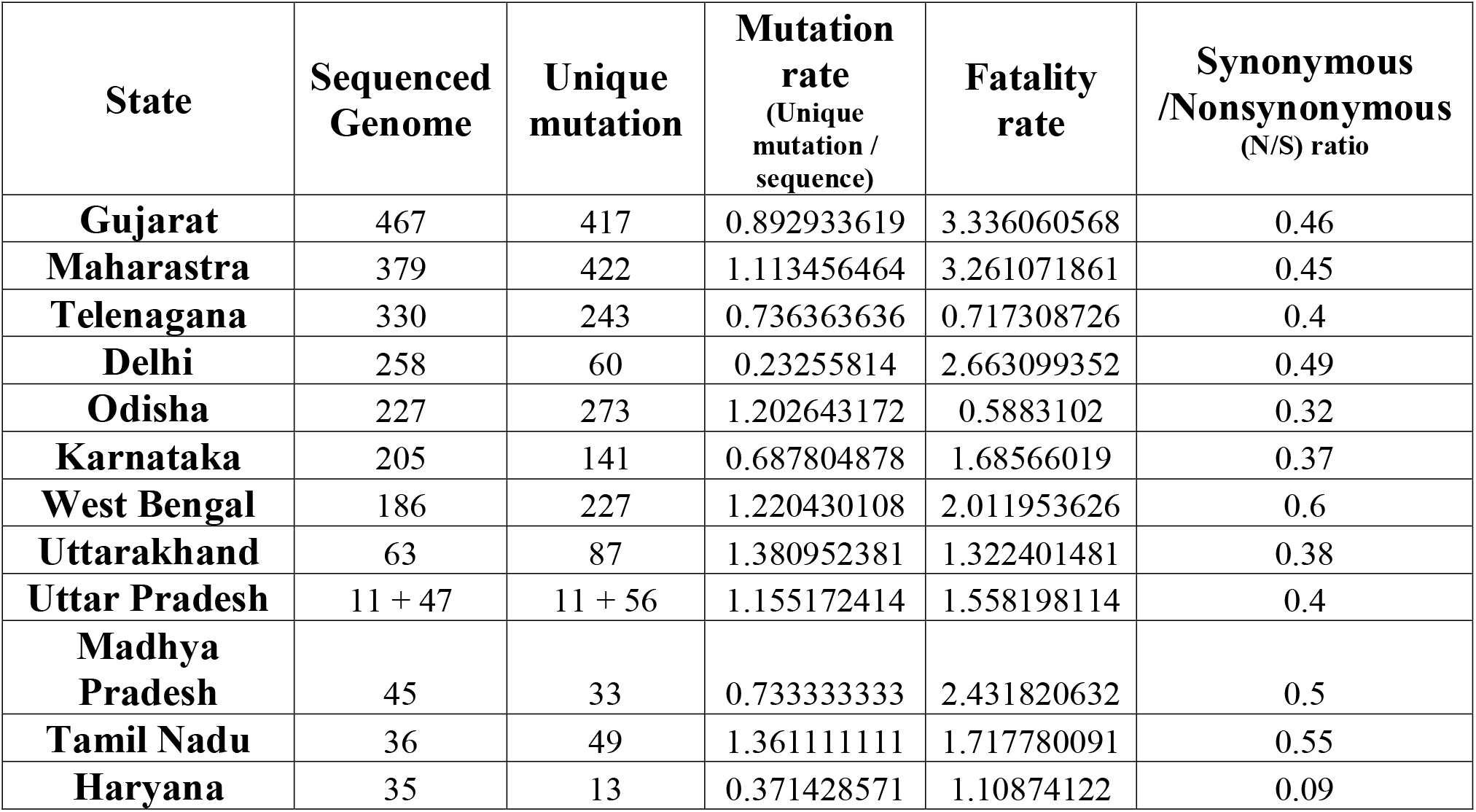
The statistics of the different selected states of India with their unique mutation rate, mortality rate and synonymous to nonsynonymous unique mutation ratio according to the mid-August, 2020 COVID-19 case.

| State | Sequenced Genome | Unique mutation | Mutation rate<br>(Unique mutation / sequence) | Fatality rate | Synonymous /Nonsynonymous (N/S) ratio |
| --- | --- | --- | --- | --- | --- |
| Gujarat | 467 | 417 | 0.892933619 | 3.336060568 | 0.46 |
| Maharastra | 379 | 422 | 1.113456464 | 3.261071861 | 0.45 |
| Telenagana | 330 | 243 | 0.736363636 | 0.717308726 | 0.4 |
| Delhi | 258 | 60 | 0.23255814 | 2.663099352 | 0.49 |
| Odisha | 227 | 273 | 1.202643172 | 0.5883102 | 0.32 |
| Karnataka | 205 | 141 | 0.687804878 | 1.68566019 | 0.37 |
| West Bengal | 186 | 227 | 1.220430108 | 2.011953626 | 0.6 |
| Uttarakhand | 63 | 87 | 1.380952381 | 1.322401481 | 0.38 |
| Uttar Pradesh | 11 + 47 | 11 + 56 | 1.155172414 | 1.558198114 | 0.4 |
| Madhya Pradesh | 45 | 33 | 0.733333333 | 2.431820632 | 0.5 |
| Tamil Nadu | 36 | 49 | 1.361111111 | 1.717780091 | 0.55 |
| Haryana | 35 | 13 | 0.371428571 | 1.10874122 | 0.09 |

**Fig. 4.**
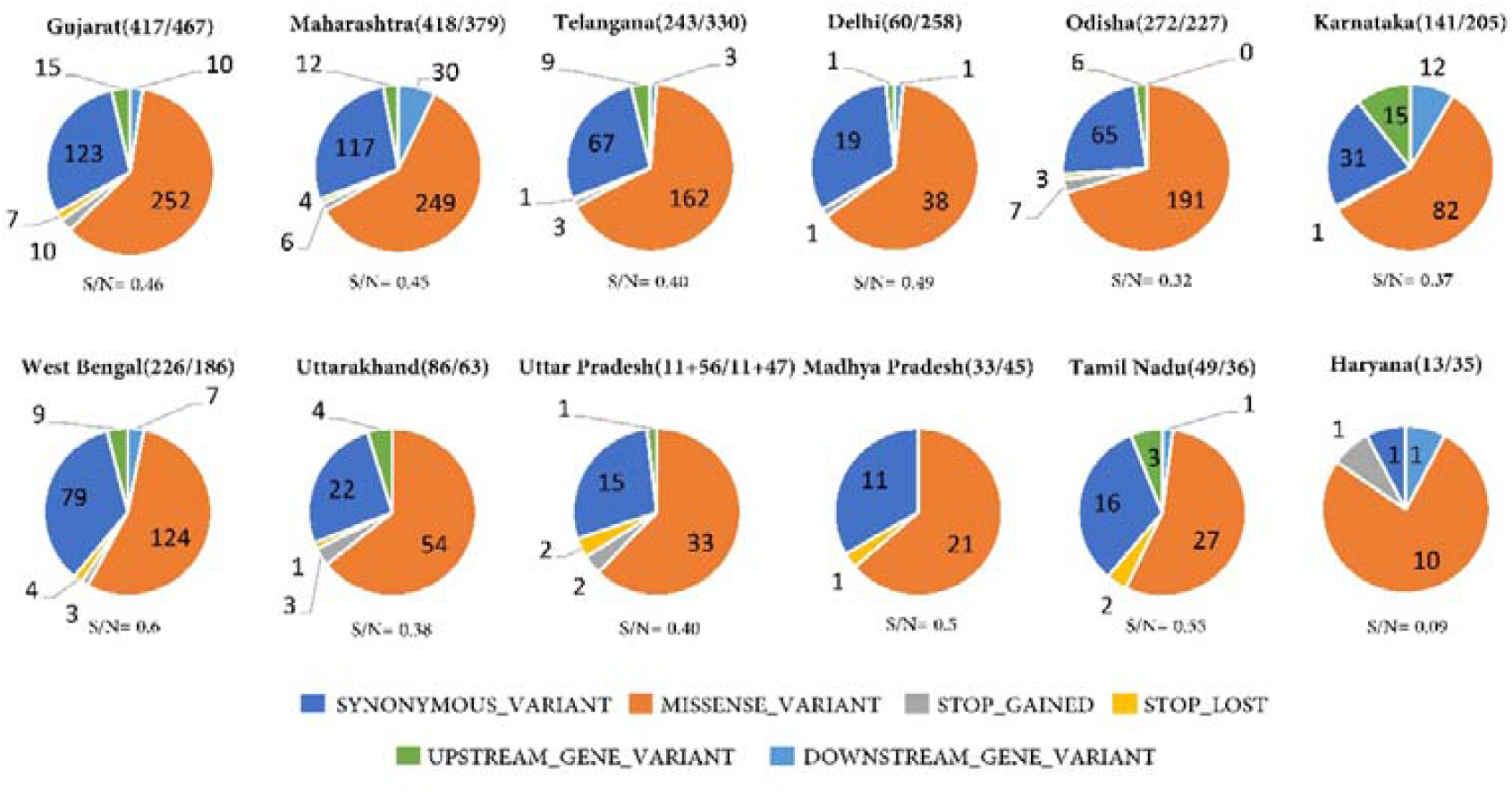
Unique mutation features in 12 different states of India whose SARS-CoV-2 genome sequences were present in GISAID database. The pie chart showed the different types of variants that were annotated on the SARS-CoV-2 genome for different states of India. The number of unique mutations variation per high-quality genome sequenced was mentioned in parenthesis. Synonymous to Nonsynonymous (S/N) unique mutation ratio of respective states were also depicted below the pie chart for each state. In the pie chart of Uttar Pradesh, the downloaded sequences and the assembled sequences were mentioned separately with plus(+) sign.

The state of Delhi had a very low unique mutation rate (0.23%) but had high death rate (2.66%). This suggests that Delhi was infected with a fatal version of the virus genome. The phylogenetic clustering by nextstrain also conformed this proposition as the genome from Delhi was clustered in 20C clade i.e. North American, which had high virulence causing large scale mortality [22]. The synonymous to nonsynonymous unique mutations (S/N) ratio of Tamil Nadu, Gujarat, Maharashtra, Delhi, Madhya Pradesh, West Bengal and Tamil Nadu was high with significantly a higher death rate (**Table 1**). Contrastingly, Karnataka, Uttar Pradesh, Uttarakhand, Haryana, Telangana and Odisha showed a relatively lesser S/N ratio (**Fig. 4**). More or less these states also showed the lower mortality rate in comparison to the overall death rate of India (1.8%). Through this observation, we can estimate the death rate by calculating the synonymous to nonsynonymous unique mutation ratio. This hypothesis could be further validated with the increasing number of SARS-CoV-2 genome sequences of all states.

### 3.6 Dynamic characterization of Unique SNP variation in UP state

We identified 56 distinct SNPs in 47 assembled sequences. Out of 56, 36 SNPs were annotated as missense variants which cause amino acid changes in the SARS-CoV-2 genome. Two nonsense mutations (T18126A, C18431T) showed the premature stop gain in the ORF1ab region (**Table 2, Fig 5**). The functional evaluation of nonsynonymous mutations predicated through Sorting Intolerant From Tolerant (SIFT) server, revealed the deleterious effects of 37% of total unique mutations. A maximum number of deleterious mutations was seen in the non-structural protein of ORF1ab region (Table 3). Threonine to Isoleucine intolerable mutation at 6056^th^ position in ORF1ab region affected the NSP11 protein in NBRI-N21 sample. This deleterious mutation might change the polymerase activity of the genomes, as it is associated with the programmed cell death evasion, cell cycle and DNA replication [35,36].

**Table 2.** Detail list of nonsynonymous SNP mutations of 47 sequences from UP with their alignment position on the genome at nucleotide and protein level. The different colored cells represent different genes while the stop codon was highlighted in green color.

| Sample_ID | POS | REF | ALT | Gene | Mutation | Type of mutation |
| --- | --- | --- | --- | --- | --- | --- |
| NBRI-N24 | 936 | C | T | ORF1ab | Thr224Ile | missense_variant |
| NBRI-N25 | 1079 | A | G | ORF1ab | Asn272Asp | missense_variant |
| NBRI-N38 | 1380 | C | T | ORF1ab | Ala372Val | missense_variant |
| NBRI-N48 | 1947 | T | C | ORF1ab | Val561Ala | missense_variant |
| NBRI-N51 | 2295 | C | T | ORF1ab | Thr677Ile | missense_variant |
| NBRI-N22 | 3741 | T | C | ORF1ab | Ile1159Thr | missense_variant |
| NBRI-N55 | 4716 | C | T | ORF1ab | Thr1484Ile | missense_variant |
| NBRI-N42 | 4973 | G | T | ORF1ab | Val1570Leu | missense_variant |
| NBRI-N47 | 5310 | C | T | ORF1ab | Thr1682Ile | missense_variant |
| NBRI-15 | 7749 | C | T | ORF1ab | Thr2495Ile | missense_variant |
| NBRI-N39 | 8359 | T | A | ORF1ab | His2698Gln | missense_variant |
| NBRI-N24 | 8636 | A | G | ORF1ab | Thr2791Ala | missense_variant |
| NBRI-N26 | 8688 | G | T | ORF1ab | Gly2808Val | missense_variant |
| NBRI-N42 | 9165 | C | T | ORF1ab | Thr2967Ile | missense_variant |
| NBRI-N44 | 12476 | A | G | ORF1ab | Met4071Val | missense_variant |
| NBRI-N30 | 14653 | G | T | ORF1ab | Met4796Ile | missense_variant |
| NBRI-N48 | 15543 | G | A | ORF1ab | Arg5093Gln | missense_variant |
| NBRI-N21 | 16912 | G | T | ORF1ab | Leu5549Phe | missense_variant |
| NBRI-N49 | 19170 | C | T | ORF1ab | Ser6302Leu | missense_variant |
| NBRI-N50 | 19374 | C | T | ORF1ab | Ser6370Phe | missense_variant |
| NBRI-N21 | 19586 | C | T | ORF1ab | Leu6441Phe | missense_variant |
| NBRI-N58 | 21911 | C | A | S | Leu117Ile | missense_variant |
| NBRI-N32 | 22436 | G | T | S | Ala292Ser | missense_variant |
| NBRI-N49 | 23126 | G | T | S | Ala522Ser | missense_variant |
| NBRI-N48 | 24620 | G | T | S | Ala1020Ser | missense_variant |
| NBRI-N35 | 24680 | G | T | S | Val1040Phe | missense_variant |
| NBRI-N48 | 25556 | T | G | ORF3a | Val55Gly | missense_variant |
| NBRI-N39 | 26006 | G | C | ORF3a | Ser205Thr | missense_variant |
| NBRI-N44 | 26158 | G | T | ORF3a | Val256Phe | missense_variant |
| NBRI-N24 | 27493 | C | T | ORF7a | Pro34Ser | missense_variant |
| NBRI-N34 | 27994 | A | G | ORF8 | Asp34Gly | missense_variant |
| NBRI-N20 | 28545 | C | T | N | Thr91Ile | missense_variant |
| NBRI-N50 | 28869 | C | T | N | Pro199Leu | missense_variant |
| NBRI-N57 | 29109 | C | A | N | Pro279Gln | missense_variant |
| NBRI-N29 | 29383 | G | T | N | Lys370Asn | missense_variant |
| NBRI-N34 | 29385 | A | T | N | Asp371Val | missense_variant |
| NBRI-N20 | 18126 | T | A | ORF1ab | Leu5954* | stop_gained |
| NBRI-N50 | 18431 | C | T | ORF1ab | Gln6056* | stop_gained |
| NBRI-N28 | 19933 | A | T | ORF1ab | Ter6556Cysext*? | stop_lost |

**Table 3.** The Putative nonsynonymous mutation with their SIFT score. The SIFT score value less than 0.05 are depicted as a deleterious mutation, highlighted in yellow whereas the unhighlighted SIFT score values are predicted as a tolerable mutation.

| Sample_ID | Gene | Mutation | SIFT Score |
| --- | --- | --- | --- |
| NBRI-N34 | ORF1ab | P5496S | 0.81 |
| NBRI-N57 |  | T6056I | 0.03 |
| NBRI-N15 |  | T2495I | 0.55 |
| NBRI-N16 |  | T2016K | 0.59 |
| NBRI-N21 |  | V4691F | 0.01 |
|  |  | V5550L | 0.02 |
|  |  | V5894I | 0.59 |
|  |  | T6441I | 0.85 |
| NBRI-N22 |  | S5483F | 0.02 |
| NBRI-N23 |  | T5300I | 0.13 |
| NBRI-N25 |  | N272D | 0.06 |
| NBRI-N27 |  | P3395S | 1 |
| NBRI-N28 |  | T6557S | 0 |
| NBRI-N30 |  | V4797F | 0 |
| NBRI-N36 |  | R24S | 0.47 |
|  |  | G7063S | 0.7 |
| NBRI-N38 |  | A372V | 0.05 |
| NBRI-N40 |  | S558F | 0.46 |
| NBRI-N42 |  | V1570L | 0.02 |
|  |  | T2967I | 0.25 |
| NBRI-N47 |  | T1682I | 0.2 |
| NBRI-N51 |  | T677I | 0.36 |
| NBRI-N55 |  | T1484I | 0.05 |
|  |  | V4980L | 0 |
| NBRI-N39 |  | H2698Q | 0.18 |
| NBRI-N24 |  | T224I | 0.03 |
| NBRI-N26 |  | G2808V | 0.14 |
| NBRI-N17 | N | D343H | 0.19 |
| NBRI-N34 |  | D371V | 0.01 |
| NBRI-N29 |  | K370N | 0.02 |
|  |  | D371Y | 0.01 |
| NBRI-N50 |  | P199L | 0.11 |
| NBRI-N57 |  | P279Q | 0 |
| NBRI-N41 |  | T362I | 0.03 |
| NBRI-N48 | S | A1020S | 0.39 |
| NBRI-N32 |  | A292S | 0.64 |
| NBRI-N35 |  | V1040F | 0.12 |
| NBRI-N49 |  | L5F, A522S | 0.10,0.73 |
| NBRI-N58 |  | L117I | 0.76 |
| NBRI-N39 | ORF3a | S205T | 0.06 |
| NBRI-N48 |  | V55G | 0 |
| NBRI-N24 | ORF7a | P34S | 0 |
| NBRI-N29 | ORF8 | D34G | 0 |
| NBRI-N26 |  | V62L | 0.54 |

**Fig. 5.**
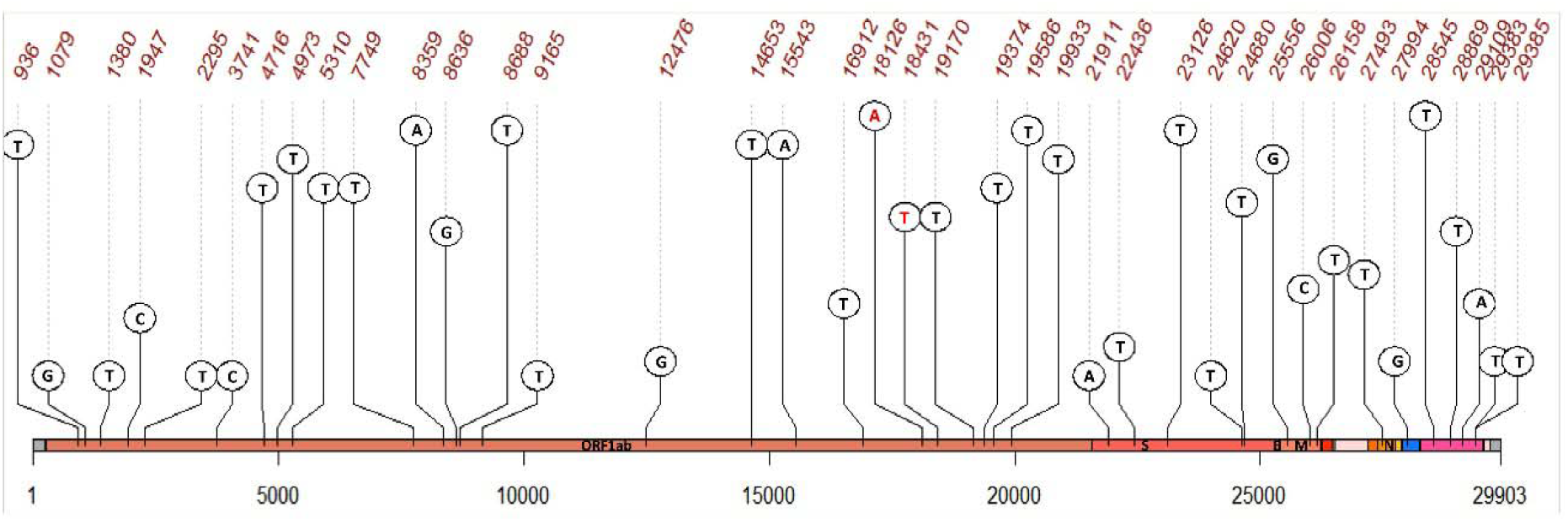
The nonsynonymous mutation variation of the SARS-CoV-2 genome for 47 sequenced samples from Uttar Pradesh. The scale bar shows the length of the genome. Dashed line with a brown color number represents the putative mutation on the genome. Black color bases are the missense variations while red color shows the stop gain at the genomic region.

One deleterious mutation was found in ORF3a, ORF7a, and ORF8 regions of the SARS-CoV-2 genome (**Table 3**). The non-structural proteins present in these ORFs act as an accessory factor [37], and assisted for virus-host interactions [38]. ORF3a regulates the interferon signaling pathway and cytokines production [39]. Deleterious mutation in the accessory protein has been reported in the latest evolving SARS-CoV-2 genome from Indian populations [40] which corroborates our finding. A structural protein, N encodes nucleocapsid protein shows the deleterious effects by the mutations at D371V, K370N, D371Y, P279Q, and T362I positions. This protein has a highly conserved domain [41] that plays a crucial function in the virus genome by regulating RNA transcription and modulating the infected cell biological processes [42]. Another nonsynonymous, deleterious mutation (V1570L) was observed in NSP3 protein in the NBRI-N42 sample. This protein has papain-like viral protease activity to generate the other replicase subunits from nsp1 to nsp16 [43].

## 4. Conclusion

In this study, 47 SARS-CoV-2 genomes from Uttar Pradesh were sequenced with high coverage third-generation sequencing technique. We identified 56 unique SNPs in our sequenced genome in which a large number of SNPs in the ORF1ab region are deleterious. These SNP variations might affect the replication of the virus genome during the infection which could be the reason for the less death rate. The identified mutations in SARS-CoV-2 genome of 47 individuals could be used as the potential target for personalized medications or effective vaccine doses to combat the effects of the COVID-19. Additionally, the relation with the synonymous to nonsynonymous unique mutation ratio with the mortality could be studied further to understand the putative region/SNPs that cause fatal to the human being.

## Author contributions

Priti Prasad: Conceptualization; Data curation; Supervision; Formal analysis; Investigation; Methodology; Resources; Software; Visualization; Writing - original draft; Writing - review & editing, Shantanu Prakash: Conceptualization; Methodology; Writing - review & editing, Mehar H. Asif: Conceptualization; Supervision; Writing - original draft; Writing - review & editing, Kishan Sahu: Resources; Writing - review & editing, Suruchi Shukla, Babita Singh, Hricha Mishra, Danish Nasar Khan, Om Prakash, MLB Bhatt: Resources, SK Barik: Conceptualization; Resources; Writing - review & editing, Samir V. Sawant: Conceptualization; Resources; Supervision; Formal analysis; Writing - review & editing, Amita Jain: Conceptualization; Resources; Writing - review & editing, Sumit Kr. Bag: Conceptualization; Resources; Supervision; Writing - review & editing

## Acknowledgments

Authors acknowledge the GISAID team and all those who submitted the genome to the GISAID database without which it’s impossible to conduct the research. PP and KS acknowledged the University of Grant Commission for providing the Senior Research fellowship. The Institute manuscript number is CSIR-NBRI_MS/2020/08/04.

